# Flexibility to contingency changes distinguishes habitual and goal-directed strategies in humans

**DOI:** 10.1101/107078

**Authors:** Julie J. Lee, Mehdi Keramati

## Abstract

Decision-making in the real world presents the challenge of requiring flexible yet prompt behavior, a balance that has been characterized in terms of a trade-off between a slower, prospective goal-directed model-based (MB) strategy and a fast, retrospective habitual model-free (MF) strategy. Theory predicts that flexibility to changes in both reward values and transition contingencies can determine the relative influence of the two systems in reinforcement learning, but few studies have manipulated the latter. Therefore, we developed a novel two-level contingency change task in which transition contingencies between states change every few trials; MB and MF control predict different responses following these contingency changes, allowing their relative influence to be inferred. Additionally, we manipulated the rate of contingency changes in order to determine whether contingency change volatility would play a role in shifting subjects between a MB and MF strategy. We found that human subjects employed a hybrid MB/MF strategy on the task, corroborating the parallel contribution of MB and MF systems in reinforcement learning. Further, subjects did not remain at one level of MB/MF behavior but rather displayed a shift towards more MB behavior over the first two blocks that was not attributable to the rate of contingency changes but rather to the extent of training. We demonstrate that flexibility to contingency changes can distinguish MB and MF strategies, with human subjects utilizing a hybrid strategy that shifts towards more MB behavior over blocks, consequently corresponding to a higher payoff.

**Author Summary:** To make good decisions, we must learn to associate actions with their true outcomes. Flexibility to changes in action/outcome relationships, therefore, is essential for optimal decision-making. For example, actions can lead to outcomes that change in value – one day, your favorite food is poorly made and thus less pleasant. Alternatively, changes can occur in terms of contingencies – ordering a dish of one kind and instead receiving another. How we respond to such changes is indicative of our decision-making strategy; habitual learners will continue to choose their favorite food even if the quality has gone down, whereas goal-directed learners will soon learn it is better to choose another dish. A popular paradigm probes the effect of value changes on decision making, but the effect of contingency changes is still unexplored. Therefore, we developed a novel task to study the latter. We find that humans used a mixed habitual/goal-directed strategy in which they became more goal-directed over the course of the task, and also earned more rewards with increasing goal-directed behavior. This shows that flexibility to contingency changes is adaptive for learning from rewards, and indicates that flexibility to contingency changes can reveal which decision-making strategy is used.

## Introduction

For optimal decision-making, animals must learn to associate the choices they make with the outcomes that arise from them. Classical learning theories suggest that this problem is addressed by habitual or goal-directed strategies for reinforcement learning [1, 2]. These strategies differ in that habitual behavior seeks simply to reinforce responses based on environmental cues, whereas goal-directed behavior considers action-outcome relationships – that is, contingencies – in the environment. Habitual and goal-directed strategies have been implemented in model-based (MB) and model-free (MF) reinforcement learning algorithms, respectively. Both algorithms make decisions by estimating action values and choosing the actions that maximize reward in the long term [3, 4]. The MF system achieves this retrospectively, caching past rewards using a reward prediction error signal [5] whereas the MB system achieves this prospectively, planning using a learned internal model of the state transitions and rewards in the environment [6,7].

Recent studies have emphasized that MB and MF systems work in parallel rather than in isolation [4, 8-10]. Early studies discerned MB and MF contributions using manipulations of reward values, such as in reward devaluation paradigms, but did not seek to quantify their relative contributions [1]. A recent study [8] addressed this by developing the hallmark “two-step” task in which each trial following rare or common outcomes was informative of the MB/MF tradeoff, thereby permitting model-fitting analyses to quantify their relative influence in decision-making. Human subjects showed a hybrid MB/MF strategy in the task, a result that has been widely replicated under different manipulations [11, 12, 13] and extended to the non-human animal literature (Groman et al. Soc. Neurosci. Abstracts 2014, 558.19, Miranda et al. Soc. Neurosci. Abstracts 2014 756.09, Akam et al. Cosyne Abstracts 2015, II-15; Hasz & Redish, Soc. Neurosci. Abstracts 2016 638.08). All these studies measured MB/MF contributions in terms of behavioral flexibility following reward updates, whereby “rare” (as opposed to “common”) observations of a rewarded or unrewarded outcome was informative to the MB system, but not the MF system, and thus these observations disclosed which system controlled participants' choices.

Theory predicts that flexibility to transition contingency changes can – like flexibility to reward structure – determine the relative influence of MB and MF strategies [4, 14]. Two studies have examined the flexibility of MB and MF systems to global contingency changes [15, 16]. However, quantification of the MB/MF tradeoff was limited as these studies manipulated contingency and tested flexibility to the change of contingency in separate phases; at these timescales, it becomes difficult to exclude the effect of adaptation on MB/MF weights. Thus, we developed a novel two-level contingency change task containing multiple, frequent and interleaved transition contingency changes that elicit different consequent actions by the MB and MF systems. Our design, like the two-step task [8] and its variants, therefore permits model-fitting analyses to robustly determine the relative influence of the MB/MF systems. The contingency change task is structured such that actions following frequent contingency changes are distinctly attributed to either a MB or MF strategy; this then permits quantification of the degree to which each system is in control.

On top of a hybrid MB/MF strategy, subjects may not remain at one level of MB/MF control but instead shift their relative weight in accordance with environmental factors. In general, animals show habit formation with time, a robust effect reported since early reward devaluation studies [17] in which extensive training stamped in habits, resulting in insensitivity to reward devaluation; in contrast, limited training retained goal-directed behavior. Sensitivity to contingency degradation (the omission of a previously-learned contingency between actions and outcomes) also decreases with overtraining, likewise reflecting a trend towards habitization with time [18]. In the original two-step task, the MB/MF trade-off was designed to be stable [8], but will shift under manipulations such as limited time [10] or cognitive load [11, 19, 20]. However, habits are not guaranteed to form with time; even after extended training, rats can show residual flexible responding following outcome devaluation, indicating that they retained goal-directed behavior despite overtraining [21]. In another study using the two-step task [20], the level of MB/MF control in fact increased in favor of more MB control (i.e. towards less habitual behavior) over three days of training. However, general shifts in MB/MF control should be disentangled from the effects of environmental volatility, which are known to affect the MB/MF balance [22]. Thus, in this study, we examined whether the MB/MF relationship is affected by environmental stability, or whether it shifts more generally over time.

We found that human subjects indeed showed a hybrid strategy in reacting to contingency changes in our task, with an increased influence of MB control over the first two blocks. However, relative MB/MF control did not significantly differ across rates of contingency changes; thus, the increase in MB control may be a more global effect of “anti-habitization” over time.

## Results

Subjects (*N*=16) performed a two-level contingency change task that consisted of 600 trials (Fig 1). Each trial began at either the first level (S0) with 50% probability, or the second level with 50% probability – 25% for each of the two states at this level (S1 or S2). If a trial started at the first level, a two-alternative choice was possible between two abstract stimuli. Each first-level action always deterministically led to the same second-level state, i.e. A1 to S1 and A2 to S2. Critically however, transitions from the second-level states to the terminal states flipped between two contingencies every 3-14 trials. Each of the two terminal states was then associated with either high or low reward, with the exact reward values drifting across trials (see Methods for details). Thus, flexibility to contingency changes was essential for maximizing reward.

**Fig 1.**
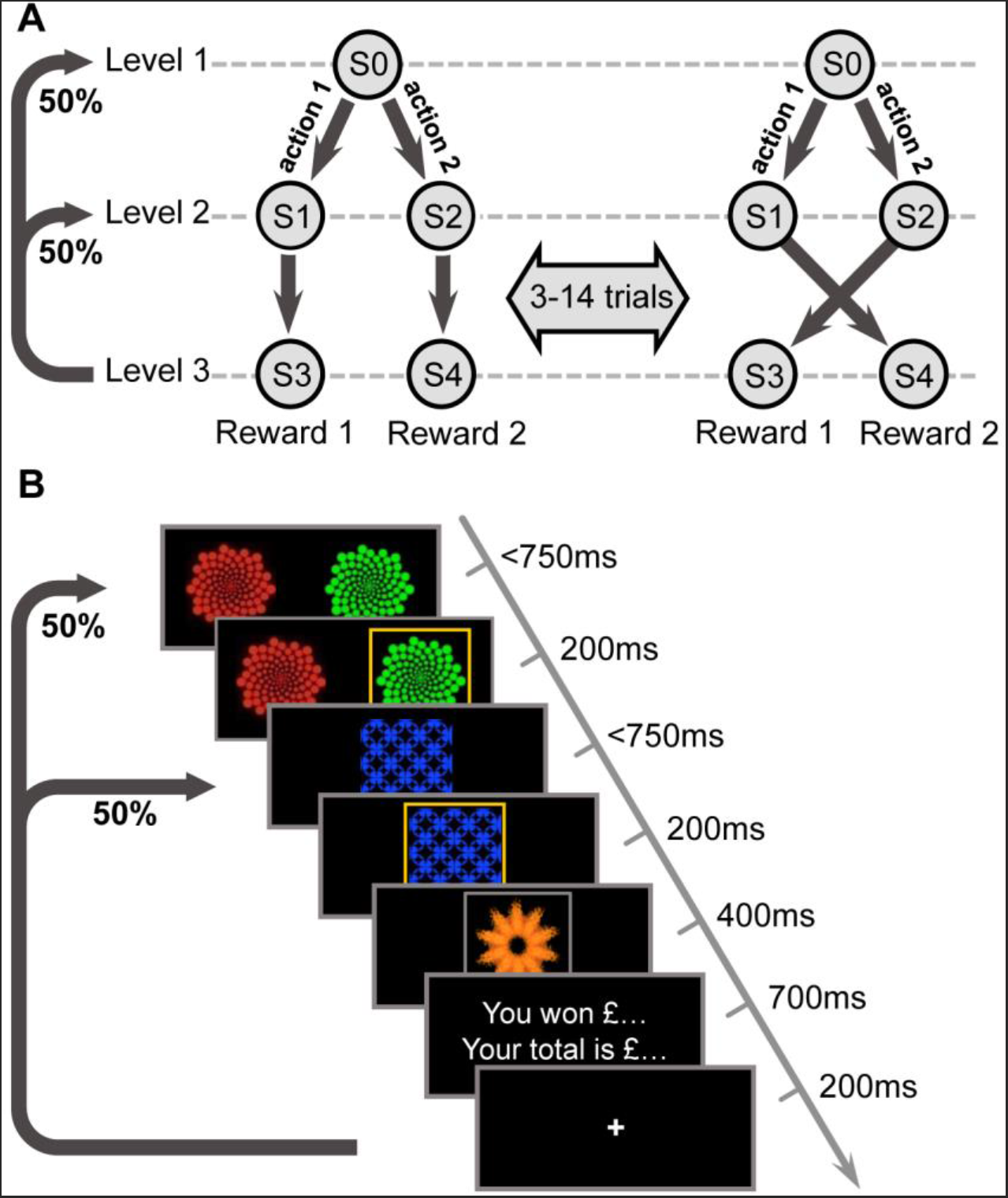
Schematic of the experimental design. (A) Each trial started from either the first-level state (S0), with 50% probability, or one of the two second-level states (S1 or S2), each with 25% probability. While two choices were available at S0, only a single forced choice was available at the second-level states. The transition structure from the second-level states to terminal states repeatedly flipped after a random number of trials (every 3-14), in an unsignalled fashion. One of the two terminal states (S3 or S4) was associated with a high reward outcome and the other resulted in a low reward outcome. (B) Timeline of the task for one example trial.

If a contingency change occurred, subjects always experienced the new transition structure regardless of whether they started at the first or second level, as contingency could only change between second-level and terminal states. Therefore, provided that an action was possible at the next trial (i.e. that the next trial started at the first level) the MB system would plan using the updated causal structure and thus would take the action that led under the new transition contingencies to the high reward terminal state. However, if a contingency change trial started from the second level, the MF system would not choose the optimal action on the next trial, as neither the received reward nor the new contingency would update the cached values of first-level actions, simply because no first-level action was experienced on those trials. As a result, the relative contribution of MB and MF systems can be measured by the degree of behavioral flexibility on first-level trials following contingency change trials starting from the second level.

To examine the effect of environmental volatility on the contribution of the two systems, the frequency of contingency changes was varied – from 3-6 trials for 200 trials, to 7-10 trials for another 200 trials, and then 11-14 trials for the final 200 trials. The order of fast and medium contingency-change blocks was counterbalanced across two subject groups (*n*=8 each). Every 40 trials, assignment of the high and low reward states also flipped to prevent formation of habits over an extended state representation, which could masquerade MF as MB behavior [23].

Simulated choices on the task were implemented according to MB and MF reinforcement learning algorithms (see Methods for details). For each system, we measured a “stay probability” index that followed the logic of contingency change trials described above. This index differs from classic stay probabilities [8] as trials starting from the second level do not have any choices to “stay”. Instead, stay probability in our task was defined as the probability of choosing the first-level action that results in the same second-level state as the previous trial. Stay probability was measured for four different conditions: whether the reward received in the previous trial was “high” or “low”, and whether the transition experienced in the previous trial, relative to the trial before that, was “changed” or remained “fixed”. Our analyses of stay/switch choices were restricted to trials that started from the first level and thus allowed the participants to make a choice. Furthermore, we restricted our analyses to first level trials following trials starting at the second level, since only these could distinguish MB and MF strategies.

Across these conditions, MB and MF systems showed different stay probability patterns. The MF system, having no experience of the action that led to the new contingency, was more likely to stay on the action leading to the high reward state, and shift on the action leading to the low reward state, under “fixed” than “changed” conditions (*p* < 0.01), indicating it was not flexible to changes in contingencies (Fig 2A). However, the MB system could immediately adapt with the correct next action, staying on the action if it would lead to the high-reward state but shifting if it would lead to the low-reward state, with a main effect of reward (*p* < 0.01) regardless of contingency condition (Fig 2B). As expected, for contingency changes from the first level, MB and MF systems did not differ in stay probability patterns, as the MF system was able to update its action values accordingly, given that it directly experienced the action leading to the new contingency (S1 and S2 Figs). In addition to pure MF and pure MB strategies, we simulated a hybrid model that linearly weights MB and MF action values according to a parameter *wMB*. The stay probability pattern produced by this hybrid system reflected a mixture of the effects observed for the pure MF and MB stay probabilities – that is, showing a main effect of reward (*p* < 0.01), but also an interaction between reward and contingency (*p* < 0.01) (Fig 2C).

**Fig 2.**
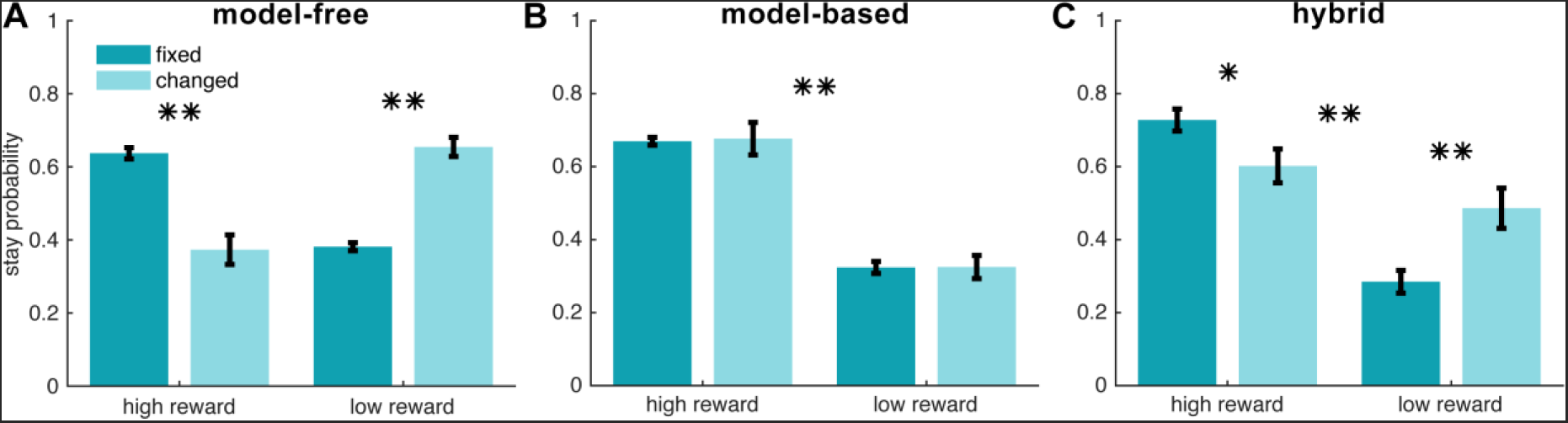
Stay probability patterns predicted by simulating model-free (A), model-based (B), and hybrid (C) reinforcement learning algorithms. Stay probability measures the probability of choosing the first-level action that results in the same second-level state as the previous trial. This index was measured for trials under four conditions: whether the reward received in the previous trial was “high” or “low”, and whether the transition experienced in the previous trial (relative to the trial before that) had its contingency “changed” or remained “fixed”. Stay probabilities are plotted for trials following a change trial that started at the second level, as only these distinguish model-free and model-based strategies. For the hybrid model (C), the parameter *wMB* was set to 0.56, the median value of the parameter inferred from participants' behavior. \**p* < 0.05, \*\**p* < 0.01

Subjects showed hallmarks of both MB and MF strategies in reacting to contingency changes (Fig 3A), showing a main effect of reward, *F*(1,60) = 24.65, *p* < 0.01, as well as a reward/contingency interaction, *F*(1,60) = 13.60, *p* < 0.01. Therefore, subjects did not solely use a MB or MF strategy when reacting to contingency changes, but rather displayed a hybrid MB/MF strategy.

**Fig 3.**
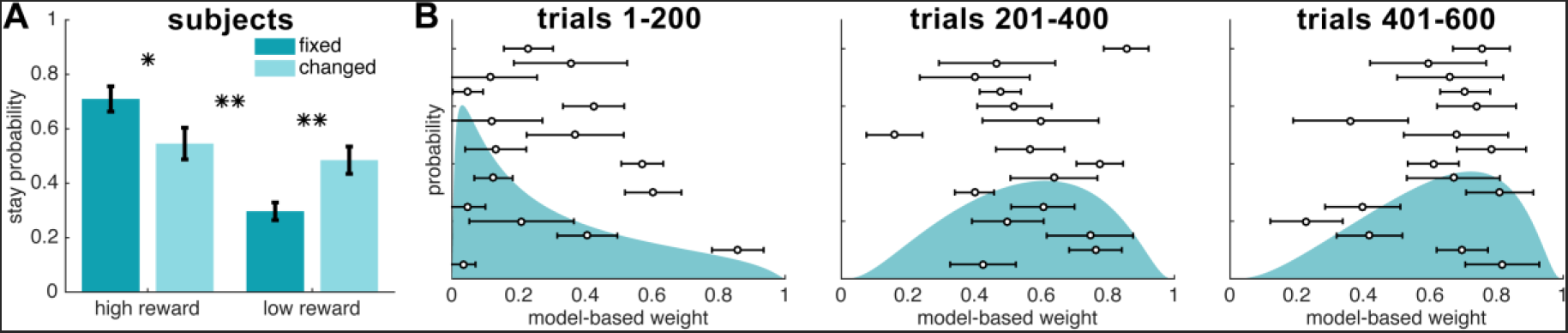
Experimental results. (A) Stay probabilities from human subjects (*N*=16) showed significant effects of both model-based (*p*<0.01) and model-free (*p*<0.01) strategies. \**p* < 0.05, \*\**p* < 0.01 (B) Probability density function over the model-based weight parameter, estimated in three different blocks of the first, middle and last 200 trials (out of 600 trials). Overlaid are the individual subjects' model-based weight parameter estimates for each block type. Error bars represent standard deviation.

While stay probabilities ruled out a purely MB or purely MF strategy, this measure could not quantify the degree to which subjects used the hybrid strategy; therefore, we used a hierarchical Bayesian method to fit candidate models of behavior to the subjects' data, to determine which model best explained subjects' choices and to obtain parameter estimates for the MB/MF weighting used by the subjects. The models tested included a pure MB model, a pure MF model, a hybrid model with one constant weight *wMB* across the session, a hybrid model with three separate *wMB* weights for each range of contingency change rates (fast, medium, or slow), and a hybrid model with three separate *wMB* weights for the three experimental blocks. The last two models served to test whether the relative contribution of the two systems depended on volatility of transition structure, or instead on block order, as contingency change rates were counter-balanced across the first two blocks. Model-fitting was confirmed to be able to recover true parameter values, as simulations showed that median estimated parameter values from model-fitting (see Methods for details) were well-correlated to known simulated parameter values, *r* ≥ 0.99, *p* < 0.01.

Model-fitting results supported the existence of a hybrid MB/MF strategy in our task. Candidate models were compared using two criteria – integrated Bayesian Information Criterion (iBIC) which controls for number of parameters [24] and exceedance probabilities [25] (S2 Table). The hybrid model with three *wMB* weights over blocks outperformed the other candidate models under both criteria, with the lowest iBIC and a probability of 89.4% that it was the most common of the four models across subjects. Thus, from here we only discuss the results of best-fit model, the three-block hybrid model.

The median fitted *wMB* weights in the three-block hybrid increased across the three blocks (Fig 3B-D), indicating some extent of “anti-habitization” rather than habit formation. The increase of *wMB* from block 1 to block 2, but not the increase from block 2 to block 3, was significant according to permutation tests, *p* < 0.01. Stay probability analyses were not conducted on the three separate blocks, as slower contingency changes meant that the later blocks had fewer samples of contingency changes for comparison. The increase in *wMB* across blocks was not attributable to differences in quality of fit from the model-fitting procedure, as the log-likelihood of parameter estimates did not differ significantly across blocks, *F*(2,45) = 1.42, *p* > 0.05. Strength of correlations between simulated *wMB* weights and *wMB* weights recovered from simulation were also similar across blocks (block 1: *r* = 0.99, block 2: *r* = 1.00, block 3: *r*= 0.99; *p* < 0.01 for all blocks). Therefore, the significant increase in *wMB* from the first to second block was not caused by differences in quality of model fit.

To confirm that the increase in model-based weight was not due to differences in the rate of contingency changes, we further analyzed the fitted weights from the three-frequency hybrid model, which had a different *wMB* assigned to each range of contingency change rates, i.e. fast (every 3-6 trials), medium (every 7-10 trials) and slow (every 11-14 trials) contingency change blocks. The estimated *wMB* weights (S3 Fig) were not significantly different between fast vs. medium, or medium vs. slow frequency of contingency change blocks in permutation tests, *p* > 0.05. Thus, the increase in *wMB* in our study seemed to be an effect of block order rather than environmental volatility from differences in contingency change rates. In summary, subjects became more model-based across the first two blocks but did not differ in MB influence between different rates of contingency changes; therefore, it seems that block order, but not contingency change volatility, affects *wMB* in our task.

As subjects became more model-based, high reward choices and consequently reward rate also increased. Choice probabilities for the high reward action differed over blocks, *F*(2,45) = 5.77, *p* < 0.01, with post-hoc tests finding a significant increase between the first and third blocks (*p* < 0.01) and the second and third blocks (*p* < 0.05). Additionally, there was a significant difference in reward rate across blocks, *F*(2,45) = 3.83, *p* < 0.05, specifically increasing between the first and third blocks (*p* < 0.05). Mean reaction time and number of missed trials due to timeout did not significantly change across blocks, *p* < 0.05; therefore, the increase in high reward choices over blocks was not necessarily because subjects were worse at the task to begin with. Two analyses were performed to rule out the possibility of practice effects driving the association between reward rate and model-based weight. Within each block, there was a significant correlation of each subject's median *wMB* and reward rate (block 1: *r* = 0.66, *p* < 0.01, block 2: *r* = 0.65, *p* < 0.01, block 3: *r* = 0.56, *p* < 0.05), indicating that on an individual subject basis, the extent of MB control was related to reward earned. Since these analyses were conducted within blocks, the association with reward rate could not be accounted for by block order.

Additionally, the hybrid model was simulated using a range of MB weights (0, 0.2, 0.4, 0.6, 0.8 and 1) using the one-weight hybrid model for simplicity. Other free parameters were set to values fitted to the participants' data. There was a significant effect of MB weight on reward rate, (*F*(5,90) = 8.5*, p* < 0.01). Therefore, MB influence in this task corresponds to a better “payoff” in terms of reward gained. However, lack of *wMB* adaptation to contingency-change rate suggests that using the cognitively-demanding MB system, at least in this task, is not motivated by its higher payoff. In other words, although being model-based does increase the payoff, it is not the *reason* for participants showing MB behavior. To further investigate this, we also computed the effect of *wMB* on reward rate, separately for different contingency change frequencies (fast/medium/slow) and found a significant effect of frequency on reward rate in a oneway ANOVA (*F*(2,45) = 5, *p* = 0.01). This shows that contingency change volatility did not affect *wMB*, despite having a significant effect on how much a MB strategy pays off. However, this absence of evidence should not be taken as evidence of absence, given the sample-size in this study. Alternatively, this absence of evidence could be because the effect of *wMB* on reward rate, though statistically significant, is in fact very small (S4 Fig). Therefore, this difference may not be apparent enough to discern or motivate a higher engagement of the MB system as contingency change frequency increases.

## Discussion

We developed a novel two-level contingency change task in which flexibility to frequently-changing transition contingencies between states could determine the extent to which subjects were using a model-based or a model-free strategy. Subjects showed a hybrid strategy when reacting to contingency changes, corroborating recent evidence of the parallel contribution of MB and MF systems in reward-guided decision-making. Importantly, this finding confirmed that changes to transition contingencies can elicit a balance of MB and MF behavior akin to changes to reward structure. Model-fitting analyses indicated that a hybrid model with three MB weights best explained subjects’ choices, with relative MB control increasing over blocks. The rate of contingency changes did not significantly shift the MB/MF balance; rather, MB control increased over the first two blocks of trials. This increase in MB control was concurrent with an increased proportion of high reward choices and consequently increased reward rate; individually, each subject’s *wMB* was also correlated with reward gained in the same block.

In all, these results illustrated that not only do subjects use a mixed MB/MF strategy, but within this hybrid strategy, the trade-off shifts towards “anti-habitization” across the first two blocks. This agrees with a previous study [20] that used the two-step task over three days, reporting that their subjects’ MB weight increased across days. One distinction between our findings is that in [20], subjects started relatively model-based (i.e. median *wMB* > 0.5) whereas in our case, subjects began relatively model-free (i.e. median *wMB* < 0.5). This difference in starting MB weight simply may be due to individual differences, which is evident even within our subject pool. Alternatively, differences could be accounted for by the relatively short reaction time limit in our task compared to theirs (750ms in ours vs. 2000ms). A shorter reaction time limit is known to provide a depth-of-planning pressure and favor more MF control [10]. Hence, our subjects may have started more model-free and only become more model-based once they mastered prospective planning of the task structure. This is supported by the lack of significant changes in reaction time across blocks, suggesting that subjects may have used the full extent of their time and eventually learned more efficient planning under time pressure, therefore showing increased MB influence over blocks.

These findings of an increase in MB control over blocks, however, goes against another study [26] using a similar task to the two-step task, that found an exponential decay in MB weight over the experimental session, or habit formation. This difference in findings is likely because they used a fixed rather than drifting amount of reward; in stationary environments such as these, habit formation can occur from overtraining, manifesting in an increase in MF rather than MB behavior [22]. Thus, these results point to the importance of maintaining a changing environment, as subjects can otherwise adapt to the change and become habitized.

Manipulations of the rate of contingency changes did not seem to affect MB/MF control. While it has been shown that environmental volatility can influence MB/MF levels in the context of common or rare updates of reward structure [22], in our case, the kind and range of contingency change volatility did not elicit a significant difference in relative MB/MF control. Further work is certainly needed to definitively rule out the possibility that environmental volatility in the form of the rates of contingency changes does not affect MB weight, but in the present study, we find that subjects did not change their use of MB control with contingency change volatility, but rather increased MB influence more generally with block order.

In conclusion, in a two-level contingency change task, subjects showed a hybrid MB/MF strategy, emphasizing their parallel contribution in reacting to changes in transition contingencies. The inclusion of multiple, frequent changes allowed us to perform model-fitting; by doing so, we found an increase in MB control over the first two blocks, a result not detectable in model-agnostic analyses alone. Our results build on the literature reporting the use of a hybrid MB/MF strategy in reacting to changes in information about reward structure, here demonstrating a mixture of strategies in reacting to multiple, frequent contingency changes that has yet been unexplored.

In addition to MB and MF systems, a third reinforcement learning algorithm known as the successor representation (SR) [15, 27] caches transitions (i.e., the probability of occupying a state after performing an action in a previous state) in a model-free fashion, but learns reward values in a model-based fashion. The SR algorithm is therefore flexible to changes in rewards (like the MB system), but inflexible to changes in contingencies (like the MF system). As a result, the MB behavior seen in the original two-step task with non-stationary reward structure [8] could be equally explained with a SR model. In other words, *wMB* and *wSR* are conflated into *wMB*. Similarly, the MF behavior in our task with non-stationary transition structure could be interpreted under a SR framework, whereby *wMF* and *wSR* are conflated into *wMF*. In this sense, the two-step task and the task presented here complement each other in providing evidence that humans use both MB and MF strategies. This novel paradigm therefore provides another avenue for exploring the relationship between MB and MF control for future studies in neuropsychiatric disorders that may differentially implicate this balance between changes in transition contingencies and changes in reward values.

## Methods

### Ethics Statement

Sixteen subjects (nine males, mean age 24 years) took part. The study was approved by the University College London Research Ethics Committee (Project ID 3450/002). All subjects provided written informed consent.

### Experimental procedure

Subjects performed 600 trials of three blocks (200 each) which differed in frequency of contingency changes: fast (contingency change every 3-6 trials), medium (every 7-10 trials) or slow (every 11-14 trials). Each subject was assigned to one of two groups (*n*=8 each), which differed by the order of presentation of fast and medium contingency change blocks, i.e. half of the subjects had fast, medium, then slow contingency changes, and the other half started with medium, fast, then slow frequency of contingency changes.

To ensure subjects understood the task structure, they were first trained with practice trials (*N*=35) which followed the same structure of the task but used practice stimuli. After this, a training session (55 trials) started which used the test stimuli but without reward. This phase was intended to introduce some familiarity to the transition relationships between states before participants were allowed to make reward-guided decisions. This was then followed by the test session using the same stimuli, but now rewarded, for 600 trials. Subjects were informed that contingency changes would occur, but did not know the frequency of changes nor that those rates would vary across the session.

At the first level, subjects had a two-alternative forced choice between two actions (pressing ‘s’ for the action available on the left side of the screen, ‘L’ for the right) with the presentation of stimuli randomized for the left/right side of the screen. To ensure that subjects recognized second-level states, they had to press ‘D’ if they encountered one of these states, and ‘K’ for the other. Both responses had a time limit of 750ms, following which the trial would end with no reward. Missed trials were not repeated.

Payoff at the high-reward terminal state varied according to a Gaussian random walk (*N*(μ = 0.5, σ = 0.2), with a drift rate of 0.15), bounded between £0 and £1. The payoff at the low-reward terminal state was £1 minus the reward of the high-reward terminal state. In practice, this resulted in an unambiguously large reward in one terminal state and an unambiguously small reward in the other terminal state. Therefore, the overall reward structure was stationary (until it flipped after every 40 trials, as noted). Subjects received a fixed proportion of their total reward gained, with payoff bounded between £5 and £25. To make the task adequately difficult and prevent formation of complex state-space representations [23], high-and low-reward assignments switched every 40 trials between the two terminal states. This change was designed never to co-occur with contingency changes.

### Model

Both model-free and model-based algorithms seek to estimate the values of state-action pairs in order to choose the actions which can maximize expected future rewards. The state space was modelled as having a first-level state *s*_0_ with two actions *a*_1_ and *a*_2_, two possible second-level states *s*_1_ and *s*_2_, and two possible terminal states *s*_3_ and *s*_4_. There was only one action available on second-level and terminal states, as the subject did not have any choices at these levels.

The model-free algorithm updates values of state-action pairs using temporal difference Q-learning [3, 28]. The reward *r*_t_ is used to compute a reward prediction error which updates action values for that state *s* and action *a* at time *t*, *Q*_*MF*_(*s*_*t*_,*a*_*t*_). At the first level *r*_*t*_ is set to be 0 as there is no reward at this level.

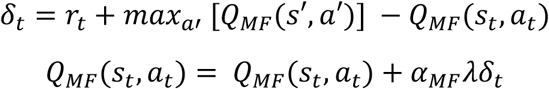

The reward prediction error updates existing action values according to a learning rate α_*MF*_, and is modified by the eligibility trace λ, which governs how much credit past actions are given for outcomes. In a *TD*(λ = 0) algorithm, first-stage actions are updated only by the second-level action values, which in turn are updated by terminal state rewards. In contrast, in a *TD* (λ = 1) algorithm, first-level actions are directly updated using the reward from the terminal state reached on that trial.

The model-based algorithm learns both transition probabilities *P*_*T*_ and reward probabilities *R*_*T*_. The transition probabilities track the transition contingencies *P*_*T*_ between states *s* and subsequent states *s*′. Upon encountering a contingency change, the model-based system always updated its knowledge of both transitions.

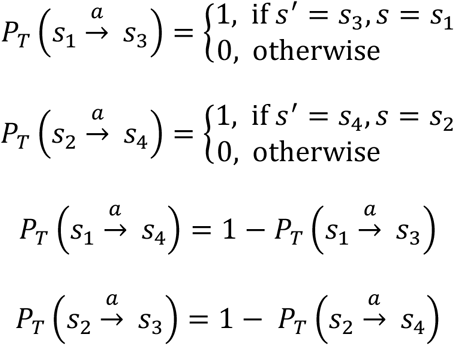

The reward probabilities *R*_*T*_ use the reward *r*_*t*_ to update its subjective reward *R* for that state *s* and action *a* at time *t*.

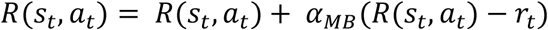

These learned transition and reward functions are then used to update the action values for the model-based system, *Q*_*MB*_.

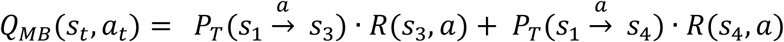

Other parameters from the simulated models included learning rates for model-based and model-free systems α_*MB*_ and α_*MF*_, and a stay bias which temporarily increased the action value for the previously-selected action regardless of outcome, to quantify a perseveration bias. These additional parameters improved fit even when controlling for model complexity (S3 Table).

For both systems, values for the non-selected action were updated as well, assuming that subjects knew that the reward for the selected action and reward for the non-selected action were negatively related, according to proposals of fictive reward [29]. Action values were updated for both visited and non-visited states, with the action values of non-visited states corresponding to 1 – *Q* (*s*_*t*_, *a*_*t*_) of the visited states. The inclusion of fictive reward updates resulted in a better fit to the subjects’ choices (S3 Table).

The hybrid model weighted MB and MF action values according to a parameter *wMB*, with *wMB*= 1 indicating fully MB control:

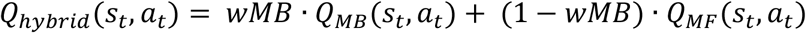

Action selection was then determined for all models according to a “softmax” rule which computes action probabilities as proportional to the exponential of the action values.

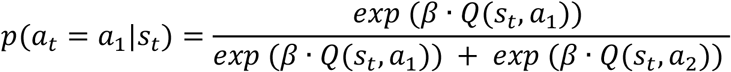

The inverse temperature *β* determined the extent to which action selection was stochastic or deterministic from action values, quantifying an exploration/exploitation trade-off.

### Simulations

To best replicate the subjects' data of 600 trials for 16 subjects, each simulation was run for 16 initializations of 600 trials each. All reported simulations used fitted parameters from the three-block hybrid model for the learning rates α_*MF*_ and α_*MB*_, inverse temperature *β*, eligibility trace λ and stay bias (S1 Table) *wMB* values were 1 for pure MB and 0 for pure MF models.

### Model-fitting

Subjects' data were fit to the models using mixed effects hierarchical model fitting. Expectation-maximisation was used which iteratively generates group-level distributions over individual subject parameter estimates, choosing the parameters that maximizes the likelihood of the data given those estimates. In each iteration, parameters were estimated by minimizing the negative log-likelihood of parameter estimates using *fminunc* in Matlab (MathWorks).

The group-level distributions over all free parameters were assumed to be Gaussian, with no constraint. To then impose sensible constraints (0 ≤ *wMB* ≤ *1;* 0 ≤ *α* ≤ *β* ≥ 0), the original free parameters were passed through a logistic function (with slope parameter equal to one) for computing *wMB* and α, and through an exponential function for computing *β*. Consequently, although the original parameters were Gaussian, the resulting parameters of the models (*wMB, α, β*) were not necessarily Gaussian (hence the skewed distributions in Fig 3). This method is preferred to imposing hard constraints on parameters because it avoids parameters hitting boundary conditions and also remains loyal to the Gaussian assumption required for hierarchical Bayesian modelling.

To ensure the efficacy of *wMB* parameter estimation for the candidate model, each block *wMB* was simulated for 11 different parameter values: 0, 0.1, 0.2,… 1. These resulted in a total of 33 parameter settings for *wMB1, wMB2, wMB3*, with 16 iterations per setting. All other parameters in the simulations were set constant as the median parameter estimates taken from the hybrid three-block model from model-fitting on the subjects' data. The same model-fitting procedure was performed on the simulated data and estimated parameter values were extracted.

The integrated Bayesian information criterion (iBIC) [24] was used to compare the fits of candidate models to the data, with lower scores indicating better fit; this criterion penalizes more complex models. Finally, Bayesian model selection [25] was used to examine the prevalence of each model in the participant population. This quantifies an exceedance probability, the probability that each model is the most common in the subject pool.

### Permutation tests

Permutation tests were run to evaluate the probability that *wMB* could differ across blocks by chance. Subjects' blocks were randomly permuted such that each “block” contained a mixture of true first, second and third blocks. Model-fitting was run on each permutation to extract parameter estimates of *wMB* for each new “block”. The probabilities *p*(*wMB_block 2_ > wMB_block 1_*), and *p*(*wMB_block 3_ > wMB_block 2_*) were then evaluated for each permutation. The occurrences of the random permutations which had a smaller *p*(*wMB_block 2_ > wMB_block 1_*), and *p*(*wMB_block 3_ > wMB_block 2_*) than the true permutation were then tallied.

Likewise, to evaluate the effect of frequency of contingency changes, permutation tests were run to compare *wMB* for fast, medium and slow contingency change blocks. Each subject was randomly assigned to one of the two groups (which differed in the order of fast and medium contingency change blocks) then *wMB* of each frequency block was computed for each permutation. Both the aforementioned one-tailed permutation test and a two-tailed Hellinger distance permutation test were used.

### Further analyses

To rule out the possibility that the effective learning rate of the MB and MF systems – rather than their fundamental differences – produced the behavior, we conducted several further analyses. A hybrid model composed of two MF systems with small (0.25) and large (0.75) learning rates could not replicate the stay probability patterns observed from subjects and the MB+MF hybrid system. Even more extreme, a hybrid model composed of two MF systems, one with a small (0.25) learning rate and no eligibility trace (λ=0), and another with a large (0.75) learning rate and with eligibility trace (λ=1) could not replicate those patterns. Furthermore, a hybrid model composed of two MB systems with small (0.25) and large (0.75) learning rates could not replicate the patterns either. When fitted to the behavioral data, all three hybrids of MF(*α*=0.25)+MF(*α*=0.75), MF(*α*=0.25,λ=0)+MF(*α*=0.75,λ=1), and MB(*α*=0.25)+MB(*α*=0.75) performed significantly worse than the real hybrid MF+MB model, according to our measures of model comparison. Together, these results rule out the possibility of just different effective learning rates of the two systems having produced the observed behavior.

We further fitted a hybrid MF+MB with a free learning rate parameter, *αMB_Transition_* for updating transitions by the MB system (rather than assuming the parameter was 1). This model, in terms of model comparison, fit the data better than pure MB and MF models, and even slightly better than the hybrid MB(*αMB_Transition_*=1)+MF model. In all previous analyses, the non-diagonal elements of the covariance matrix were set to zero (i.e., assuming no correlation between free parameters). However, for the hybrid MB+MF with a free *αMB_Transition_*, we observed strong correlations between parameters when the covariates were allowed to change freely. The *αMB_Transition_* parameter was highly negatively correlated with *α*MF, and positively correlated with stay-bias, *wMB*(block 1), and *wMB*(block 2). We therefore decided not to rely on this model and instead, for the sake of parsimony, to use the original hybrid MB(*αMB_Transition_*=1)+MF model throughout the paper.

## Acknowledgements

Thanks to Peter Dayan for helpful discussions and comments, and Thomas Akam for comments on the manuscript.

## Supporting Information

**S1 Fig.**
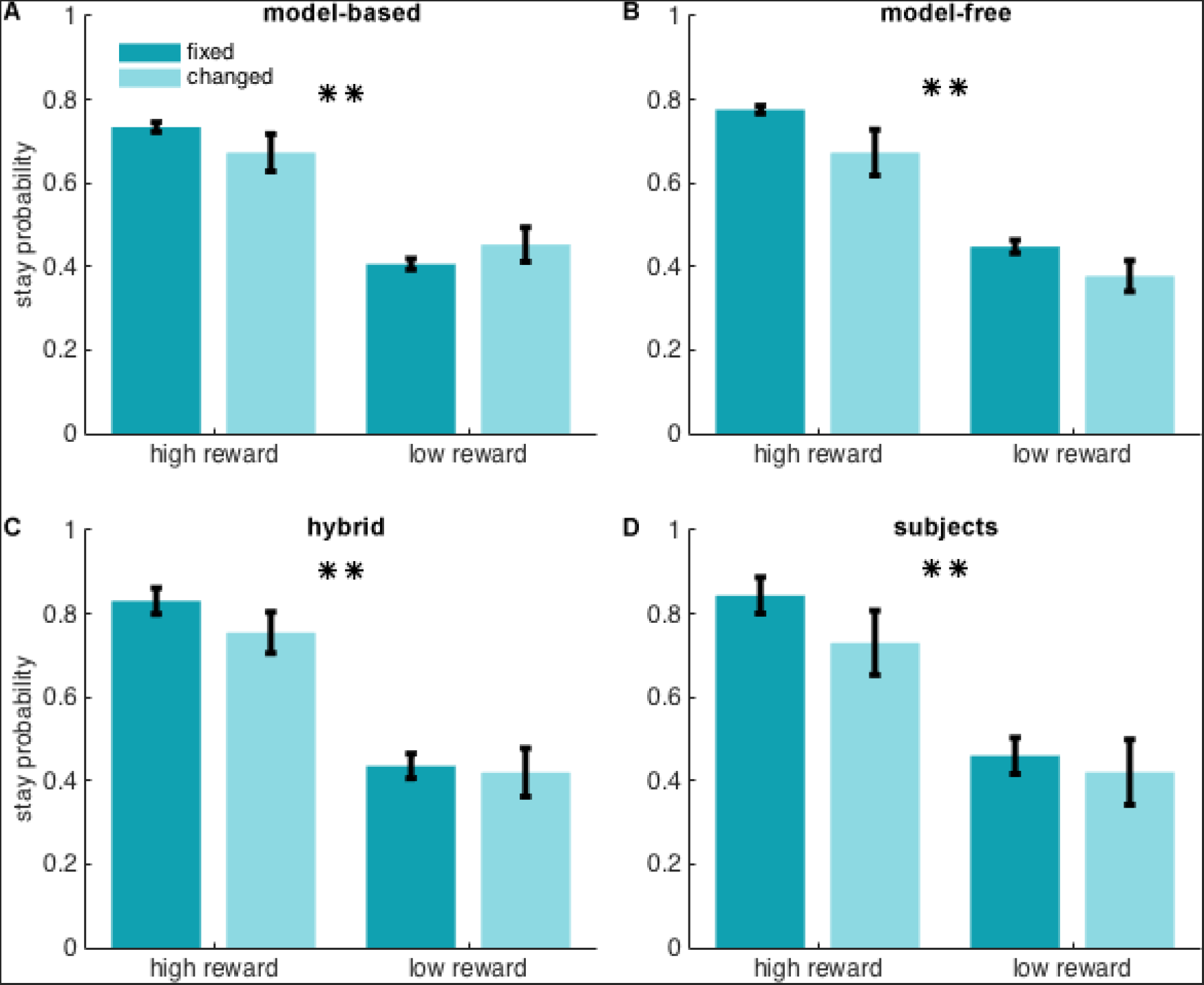
Stay probability patterns after first-level contingency changes predicted by simulating model-based (A), model-free (B), and hybrid (C) reinforcement learning algorithms, along with experimental results (D). Stay-probability measures the probability of choosing the first-level action that results in the same second-level state as the previous trial, following a trial that started at the first level. For each system, this index was measured under four different conditions: when the reward received in the previous trial was “high” or “low”, and when the transition experienced in the previous trial (relative to the trial before that) “changed” or remained “fixed”. \**p* < 0.05, \*\**p* < 0.01

**S2 Fig.**
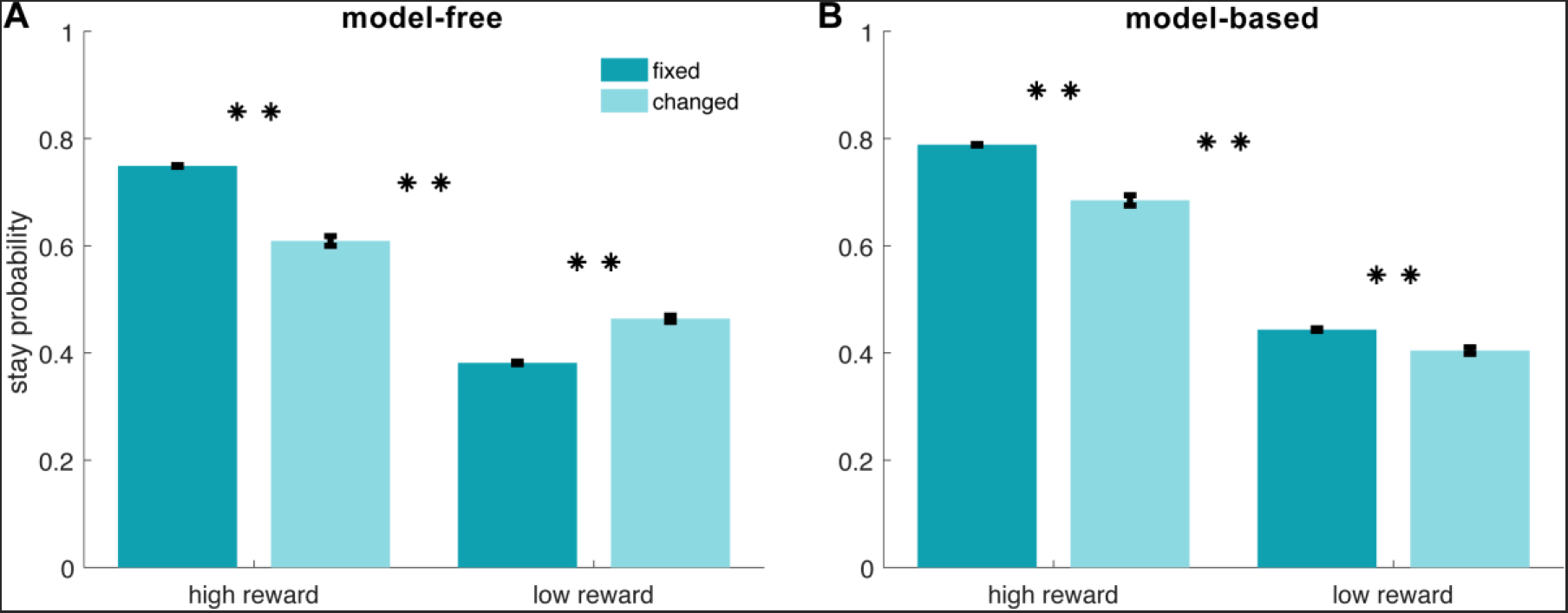
Stay probability patterns after first-level contingency changes predicted by simulating model-free (A) and model-based (B) algorithms for 1000 agents. When starting from the first level and encountering a change in the transition structure, both MB and MF systems are able to update their action values. However, the extent of this update is not equal for the two systems due to their different effective learning rates. As a result, the two systems show slightly different flexibility levels (i.e., stay-probability patterns) even when both systems were informed of the change in contingencies. Therefore, in addition to structural differences, the MB and MF algorithms that participants used in this task also have different effective learning rates. This difference is not apparent in S1 Fig because the models were only simulated 16 times (equal to the number of participants).

**S3 Fig.**
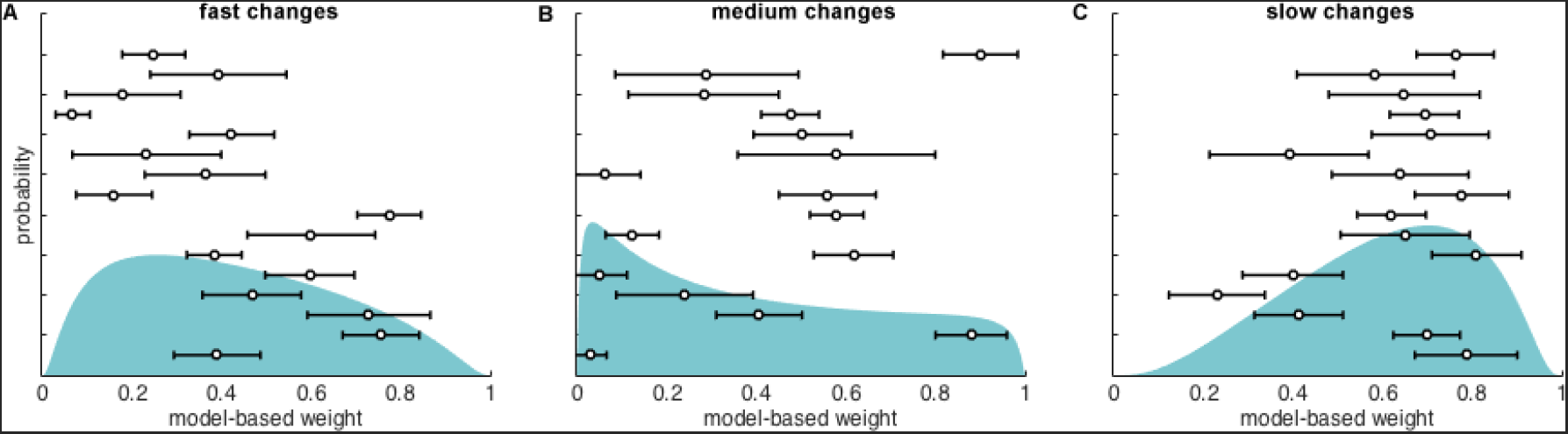
Model-based weights for (A) fast, (B) medium and (C) slow contingency changes. Probability density function over the model-based weight parameters estimated from model-fitting, for the blocks of fast (every 3-6 trials), medium (every 7-10 trials) and slow (every 11-14 trials) frequency of contingency changes. Overlaid are the individual subjects' parameter estimates for each block type. Error bars represent standard deviation.

**S4 Fig.**
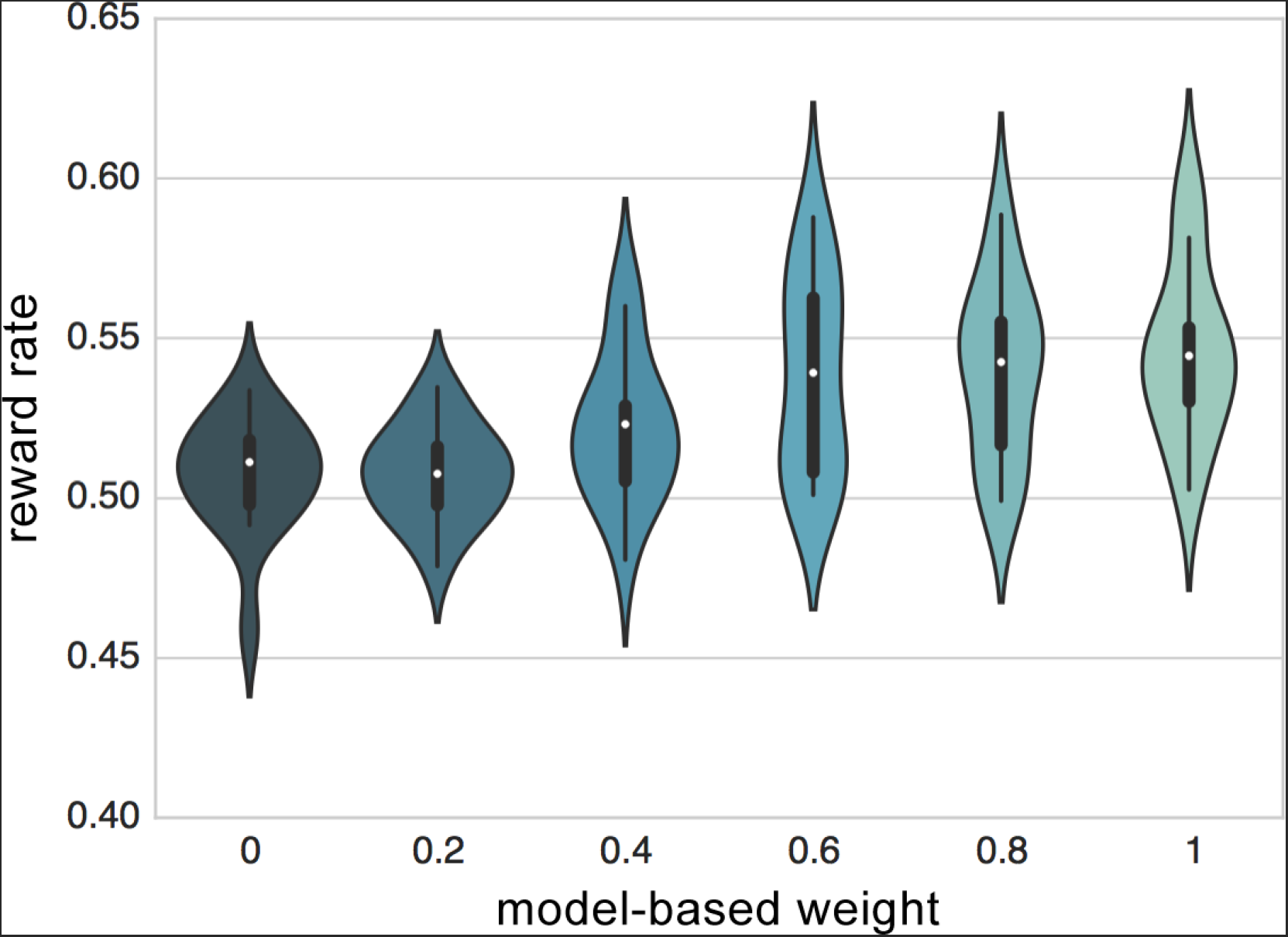
Reward rates of simulated model-based weights. Choices were simulated with six different model-based weights (0, 0.2, 0.4, 0.6, 0.8, 1, with *n*=16 iterations each) and the mean reward rate was computed. There was a significant difference in reward rate across different *wMB* values, *F*(5,90) = 8.5, *p* < 0.01, however, the difference was small, which may account for the absence of a significant modulation in *wMB* across contingency change frequencies.

**S1 Table.** Median Plus Quartile Group-level Parameter Estimates. Best-fitting parameter estimates over the subjects from model-fitting.

| | $\beta$ | Stay bias | $\alpha_{MB}$ | $\alpha_{MF}$ | $\lambda$ | $wMB$<br>(block 1) | $wMB$<br>(block 2) | $wMB$<br>(block 3) |
| --- | --- | --- | --- | --- | --- | --- | --- | --- |
| **1st quartile** | 1.88 | 0.04 | 0.55 | 0.03 | 0.30 | 0.10 | 0.40 | 0.48 |
| **Median** | 2.99 | 0.10 | 0.70 | 0.30 | 0.47 | 0.23 | 0.57 | 0.63 |
| **3rd quartile** | 4.73 | 0.22 | 0.81 | 0.85 | 0.65 | 0.46 | 0.71 | 0.76 |

**S2 Table.** Model Comparison of Candidate Models. Integrated Bayesian Information Criterion (iBIC) and negative log-likelihood of all candidate models from model-fitting. The models tested were: pure model-free (“MF”), pure model-based (“MB”), hybrid MB/MF (“hybrid”), hybrid MB/MF with different weights fitted for each of the three 200-trial blocks (“three-block hybrid”), and a hybrid model with different weights fitted for each frequency of contingency changes (“three-frequency hybrid”). The winning model was the three-block hybrid, highlighted in gray, according to iBIC [24] and Bayesian model selection [25].

| **Model** | **Model-free** | **Model-based** | **Hybrid** | **Three-block hybrid** | **Three-frequency hybrid** |
| --- | --- | --- | --- | --- | --- |
| **Parameters** | 4 | 3 | 6 | 8 | 8 |
| **iBIC** | 9051 | 8091 | 7938 | 7687 | 7711 |
| **Negative Log Likelihood** | 4489 | 4018 | 3914 | 3770 | 3782 |

**S3 Table.**
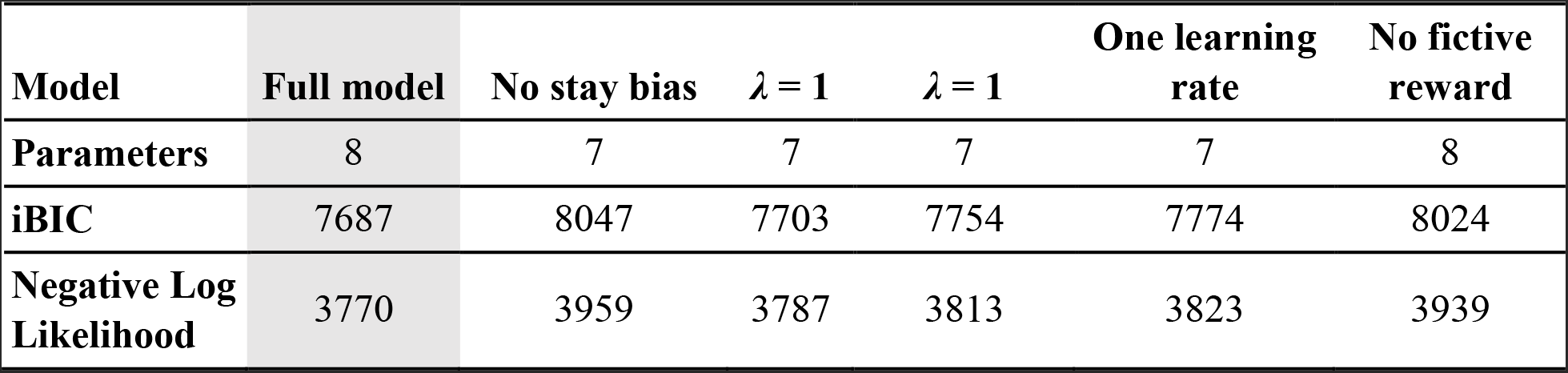
Model Comparison of Additional Parameters. Integrated Bayesian Information Criterion (iBIC) and negative log-likelihood of the winning three-block hybrid model with different weights fitted for each of the three 200-trial blocks and the same model without stay bias, with λ = 1, with λ = 0, with only one learning rate for both MF and MB systems, and without updating fictive reward. The full model fit better to the data than the same model without each of the aforementioned parameters, even when controlling for model complexity in the iBIC.

